# Explicit and implicit locomotor learning in individuals with chronic hemiparetic stroke

**DOI:** 10.1101/2024.02.04.578807

**Authors:** Jonathan M. Wood, Elizabeth Thompson, Henry Wright, Liam Festa, Susanne M. Morton, Darcy S. Reisman, Hyosub E. Kim

## Abstract

Motor learning involves both explicit and implicit processes that are fundamental for acquiring and adapting complex motor skills. However, stroke may damage the neural substrates underlying explicit and/or implicit learning, leading to deficits in overall motor performance. While both learning processes are typically used in concert in daily life and rehabilitation, no gait studies have determined how these processes function together after stroke when tested during a task that elicits dissociable contributions from both. Here, we compared explicit and implicit locomotor learning in individuals with chronic stroke to age- and sex-matched neurologically intact controls. We assessed implicit learning using split-belt adaptation (where two treadmill belts move at different speeds). We assessed explicit learning (i.e., strategy-use) using visual feedback during split-belt walking to help individuals explicitly correct for step length errors created by the split-belts. The removal of visual feedback after the first 40 strides of split-belt walking, combined with task instructions, minimized contributions from explicit learning for the remainder of the task. We utilized a multi-rate state-space model to characterize individual explicit and implicit process contributions to overall behavioral change. The computational and behavioral analyses revealed that, compared to controls, individuals with chronic stroke demonstrated deficits in both explicit and implicit contributions to locomotor learning, a result that runs counter to prior work testing each process individually during gait. Since post-stroke locomotor rehabilitation involves interventions that rely on both explicit and implicit motor learning, future work should determine how locomotor rehabilitation interventions can be structured to optimize overall motor learning.

**New and noteworthy:** Motor learning involves both implicit and explicit processes, the underlying neural substrates of which could be damaged by after stroke. While both learning processes are typically used in concert in daily life and rehabilitation, no gait studies have determined how these processes function together after stroke. Using a locomotor task that elicits dissociable contributions from both processes and computational modeling, we found evidence that chronic stroke causes deficits in both explicit and implicit locomotor learning.

## Introduction

Motor learning, the ability to acquire and maintain motor skills with practice (Schmidt and Lee 2005), involves both explicit and implicit processes. Explicit learning, as in strategy formation and use, is critical for skill acquisition because it provides the means for fast, flexible changes in movements (Bond and Taylor 2015; Huberdeau et al. 2019; Roemmich and Bastian 2018). Implicit processes keep movements finely calibrated in the face of changes to the body or environment (Bond and Taylor 2015; Krakauer et al. 2019; Mazzoni and Krakauer 2006; Roemmich and Bastian 2018). While both explicit and implicit processes typically work in concert when learning complex motor skills, stroke may damage the neural substrates underlying either component (or both), leading to deficits in overall motor learning. Since motor learning is foundational to rehabilitation of individuals post-stroke, and since explicit and implicit learning are often used concurrently (Roemmich et al. 2016; Taylor and Ivry 2011, 2012), it is important to determine if individuals post-stroke have impairments in one or both processes when assessed during the same motor task.

Explicit learning, sometimes referred to as “voluntary correction” (Roemmich et al. 2016) or “cognitive strategies” (Taylor and Ivry 2011), involves consciously directing specific changes in movement patterns. For example, a patient may consciously increase their step length in response to a clinician’s verbal instructions. Explicit learning plays a critical role in skill acquisition and motor memory for neurologically intact individuals, and is hypothesized to be mediated primarily in prefrontal cortex (Krakauer et al. 2019; McDougle et al. 2016; Taylor and Ivry 2014). We assume this process is driven by target error, the difference between a movement outcome and the task goal (French et al. 2018; Mazzoni and Krakauer 2006; Taylor and Ivry 2012). A key feature of this process is that it can be volitionally “switched” on or off in response to context or instructions (Bond and Taylor 2015; French et al. 2018; Wood et al. 2020, 2021). Explicit learning can be used during gait in both neurologically intact individuals and those post-stroke by providing visual feedback and specific task instructions (French et al. 2021b, 2021a; Hussain et al. 2013). While our prior work testing explicit learning post-stroke did not observe group-level impairments (French et al. 2021b), we did find an effect of fluid cognition on explicit learning in a separate study using the same paradigm (French et al. 2021a). This, in combination with the use of a relatively small, visual perturbation in these studies suggests the potential to observe explicit learning deficits under more challenging task conditions.

Sensorimotor adaptation is an implicit motor learning process that is essential for maintaining well-calibrated movements in response to ever-changing environments and body states. Sensorimotor adaptation, which we refer to here as “implicit adaptation”, is driven by sensory prediction error, the difference between the actual and expected sensory consequences of a motor command, and mediated in large part by the cerebellum (Krakauer et al. 2000; Morton and Bastian 2006; Shadmehr et al. 2010; Shadmehr and Mussa-Ivaldi 1994; Tseng et al. 2007). During gait, implicit adaptation can be elicited using a split-belt treadmill where the belts under each limb move at different speeds (Dietz et al. 1994). This perturbation initially produces asymmetric gait patterns (e.g., step length asymmetries) which are slowly recalibrated back to baseline asymmetry levels (Malone et al. 2012; Reisman et al. 2005). The hallmark of implicit adaptation is the storage of the adapted stepping pattern when the belts return to the same speed, termed an “implicit aftereffect” (Reisman et al. 2005).

When tested in isolation during locomotion, individuals with non-cerebellar stroke adapt their step length asymmetry to a similar magnitude as neurologically intact participants by the end of learning (Malone and Bastian 2014; Reisman et al. 2007; Savin et al. 2013; Tyrell et al. 2014) and demonstrate similar implicit aftereffects (Reisman et al. 2007; Savin et al. 2013; c.f. Malone and Bastian 2014). However, individuals with stroke adapted at a slower rate compared to controls (Malone and Bastian 2014; Reisman et al. 2007; Savin et al. 2013; Tyrell et al. 2014).

While explicit learning and implicit adaptation are typically used simultaneously to learn new skills in everyday life, including rehabilitation practice, they are mostly studied individually during gait (French et al. 2018; Hussain et al. 2013; Reisman et al. 2005; Wood et al. 2020, 2021). This may be because when they are studied within the same task, it can be difficult to dissociate the individual contributions of each process to overall behavior (Cherry-Allen et al. 2018; Long et al. 2016; Malone and Bastian 2010; Roemmich et al. 2016). A study in young neurotypical adults accomplished this using visual feedback to induce explicit learning that helped correct the step length errors produced by the split-belt treadmill (Roemmich et al. 2016). They found explicit learning improved performance during split-belt walking compared to a group that did not receive feedback. However, the implicit aftereffects (measured without visual feedback) were similar between groups, indicating explicit learning did not impact the recalibration of motor commands (i.e., implicit adaptation). Thus, the authors concluded that, within the same locomotor learning task, while explicit learning improves overall performance, implicit adaptation proceeds despite involvement from explicit learning in individuals with intact neurologic systems. Critically, it is unclear to what degree explicit learning versus implicit adaptation is impaired in individuals post-stroke when assessed in a task requiring dissociable contributions from both.

While explicit learning and implicit adaptation are broadly intact after stroke when assessed individually during gait, it is unknown if this holds when they are combined in the same task. Only two studies, both in reaching movements, have attempted to tackle this question, but with mixed results (Binyamin-Netser et al. 2023; Taylor and Ivry 2014), indicating more work is necessary to determine if there are deficits in these learning processes when assessed within the same task. This is a critical gap because gait rehabilitation post-stroke involves a combination of explicit learning and implicit adaptation (e.g., a patient may explicitly try to increase step length based on their therapist’s instructions while simultaneously implicitly adapting to a more compliant walking surface). Therefore, determining how each is impaired, when occurring together in the same task, has important implications for how rehabilitation of locomotor tasks should be optimally structured.

The purpose of this study was to determine if individuals with chronic, hemiparetic stroke demonstrate impaired explicit learning and/or implicit adaptation during a locomotor task involving dissociable contributions from both processes. We accomplished this through a combination of behavioral testing and computational modeling. Since cortical structures, and in particular, prefrontal regions contribute to cognitive processes that are often impaired in stroke (Barker-Collo and Feigin 2006), we hypothesized that individuals with stroke would demonstrate impaired explicit learning compared to controls. Additionally, because the rate of implicit adaptation on the split-belt treadmill is slow but the magnitude is intact in persons post-stroke (Reisman et al. 2007; Savin et al. 2013; Tyrell et al. 2014) we hypothesized that individuals with stroke would demonstrate similar levels of implicit adaptation as controls.

## Materials and methods

### Participants

We recruited 21 (10 Female) individuals with one prior unilateral, stroke to participate in this study and 18 (9 Female) healthy age- and sex-matched control participants. Individuals with stroke were included if they were between 18 and 85 years old, had a single unilateral hemiparetic stroke (confirmed by an MRI or CT scan) more than 6 months prior, and were able to walk without assistance from another person. Individuals with stroke were excluded if they had evidence of cerebellar stroke, other neurologic diagnoses aside from stroke, inability to walk outside of the home prior to stroke, pain limiting walking, neglect, or significant aphasia. Control participants were excluded if they had any conditions that might limit their walking or motor learning, any neurologic conditions, or uncorrected vision or hearing loss. All individuals provided written informed consent prior to participating and the study was approved by the University of Delaware Institutional Review Board.

### Experimental design

To determine if individuals with stroke have impaired explicit learning or implicit adaptation during a locomotor learning task that requires contributions from both processes, we combined the split-belt adaptation paradigm with real-time visual feedback, similar to a previous study (Figure 1A; Roemmich et al. 2016). Participants performed 4 phases of treadmill walking: Baseline, Practice, Adaptation, and De-adaptation (Figure 1B). During Baseline and Practice, both the treadmill belts moved at the same speed, set at half the speed of the fast belt. During the Baseline phase, no visual feedback was provided on the screen and individuals were told to “walk comfortably”. The Practice phase served to introduce participants to the visual feedback and ensure the individuals with stroke could respond to the visual feedback by changing their step length. Step length targets first appeared at each participant’s baseline step length for 90 seconds, at which point, they were verbally oriented to the feedback and instructed to practice changing their step lengths by stepping both above and below the targets. For the next 30 seconds, the step length targets shifted 10 cm longer for the limb taking the longer baseline step and 10 cm shorter for the limb taking the shorter baseline step. The targets shifted back to the baseline step lengths for the final 60 seconds of the Practice phase and individuals were asked to “walk comfortably”. We confirmed that participants in the stroke and control groups could change their step lengths from baseline in response to the visual feedback (step length change from targets shift to walk comfortably, stroke fast = 4.6 cm SD 3.6, slow = -3.0 SD 3.5; control fast = 7.8 SD 3.5, slow = -7.2 SD 3.2).

**Figure 1.**
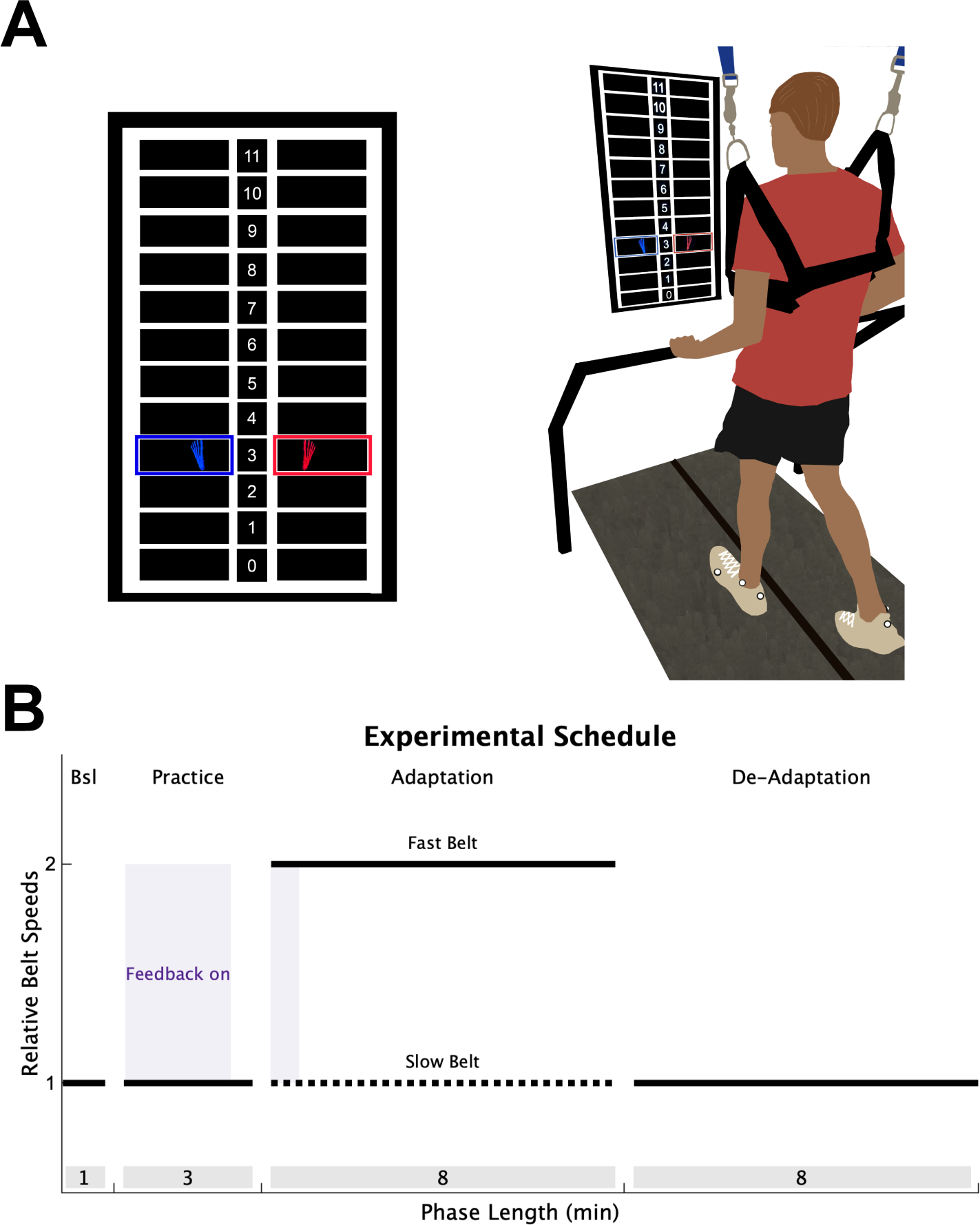
Experimental Design. (**A**) Individuals walked on a split-belt treadmill with a vertically mounted television screen in front of them. The visual feedback was a grid of 12 different step lengths, each 10 cm in height. The step length feedback was represented on the screen as blue (left) and red (right) feet that appeared on the screen as soon as heel strike was detected and disappeared once toe off was detected. **(B)** All participants completed 4 walking phases: 1) A Baseline (Bsl) phase of normal walking where no feedback was on the screen; 2) A Practice phase where individuals were introduced to the visual feedback while walking (purple shading); 3) An Adaptation phase where the slow belt (dotted black line) moved at half the speed of the fast belt (solid black line), with feedback activated during only the first 40 strides (purple shading); 4) A De-adaptation phase where the belts returned to the same speed. The length of each phase (in minutes) is displayed in the grey shading at the bottom of the figure.

During the Adaptation phase (8 minutes), the fast belt speed was set at the fastest overground gait speed (stroke group mean = 1.29 m/s SD 0.38; control = 1.81 SD 0.22), and the slow belt moved at half the speed of the fast belt, producing a 2:1 speed ratio (Charalambous et al. 2018; Tyrell et al. 2014). To ensure there were not large differences in treadmill speeds between the stroke and control groups, we constrained the fast treadmill belt speed between 0.6 and 1.0 m/s for all participants. This constraint resulted in both groups walking at similar speeds (stroke group mean = 0.94 m/s SD 0.12; control = 1.0 SD 0). For participants in both groups, the limb that took the longer step during the Baseline phase was placed on the fast belt. This perturbation produces a large asymmetry of the left and right step lengths (defined as the distance between two feet at heel strike), and is corrected on a stride-by-stride basis through implicit adaptation (Charalambous et al. 2018; Reisman et al. 2005, 2007). Lastly, participants performed a De-adaptation phase (8 minutes) where they were instructed to “walk comfortably”, and both belts moved at the same speed as the Baseline phase (i.e., the slow belt speed) so that we could measure the size of the implicit aftereffect, our measure of the total magnitude of implicit adaptation.

To assess explicit learning, defined as the ability to consciously correct for errors between a movement outcome and the task goal, we provided visual feedback of the left and right step lengths during the first 40 strides of the Adaptation phase. The real-time visual feedback was displayed on a vertically orientated LCD television screen placed 100 cm in front of the treadmill (Figure 1A; Size: 123.3 x 71.1 cm; Sony Tokyo, Japan). The Motion Monitor software (Innovative Sports Training Inc., Chicago, IL, USA) was used to display the visual feedback during the experiment. The feedback consisted of a target grid of 12 possible step lengths, each 10 cm in height. This grid had a 1:1 correspondence with the actual step length. The left and right step length feedback was displayed as a red and blue foot, respectively. Each foot was presented in the center of the row corresponding to that step length window, and appeared as soon as heel strike was detected, then disappeared once the subsequent swing phase began. The target right and left step lengths during the Adaptation phase were set at each participant’s left and right baseline step lengths, denoted by highlighting the corresponding row of the grid. Participants were instructed to “hit the targets” when the feedback was visible. Therefore, because the targets were set at baseline step length, the feedback guided participants to voluntarily correct the step length asymmetry induced by the split-belt treadmill via explicit learning. We chose these targets to prevent any ambiguity in the explicit task goal, and we reasoned that baseline step lengths were most parsimonious targets to use.

The key manipulation that allowed us to assess the magnitude of explicit learning was to turn off the feedback after the first 40 strides of Adaptation and instruct participants to “walk comfortably”. Here, we relied on the flexibility of explicit learning, assuming it could be voluntarily “switched off”, leaving no residual aftereffects due to implicit adaptation (Bond and Taylor 2015; McDougle and Taylor 2019; Wood et al. 2020). Meaning, after the instructions and removal of feedback, only implicit adaptation remained. Thus, our measure of the total magnitude of explicit learning was the difference between the step length behavior when the feedback was on and when it was first turned off (Roemmich et al. 2016).

### Data collection

During all phases of treadmill walking, individuals wore a ceiling mounted harness (that did not provide body weight support) and held a handrail to prevent falls. Additionally, we monitored heart rate (Polar, Kempele, Finland) for safety, and determined perceived exertion using the Borg Rate of Perceived Exertion (RPE) scale after each walking phase. If participants exceeded >80% their age-predicted max heart rate, the treadmill was stopped, and the participant was provided with a seated rest break until their heart rate recovered.

However, no included participants needed the treadmill to stop during any of the walking phases.

Participants walked on a dual belt treadmill that captured kinetic data through two force plates, one under each belt, at 1000 Hz (Bertec, Columbus, OH, USA). Kinematic data were captured and recorded at 100 Hz using a Vicon MX40, 8-camera motion capture system, and time-synchronized with the kinetic data in Nexus software (v2.8.2, Vicon Motion Systems, Inc., London, UK). We used a custom marker set with seven retroreflective markers, one for each heel, lateral malleolus, and fifth metatarsal head, and the left medial malleolus.

### Data analysis

Step length, calculated in real-time using motion capture and the Motion Monitor, was defined as the anterior-posterior distance between the two ankle markers at heel strike. Heel strike was determined in real time using the following criteria: 1) The leading limb’s heel marker must be anterior with respect to the trailing limb’s heel marker; 2) a ground reaction force > one-third of the participants body weight detected through the treadmill force plate; and 3) no ground reaction force > one-third of the participants body weight detected through that same belt 40 ms prior to the detected step length.

The remainder of the data were analyzed with custom written MATLAB scripts (vR2022a, Mathworks, Natick, MA, USA). Step lengths were used to calculate step length asymmetry on each stride (s):

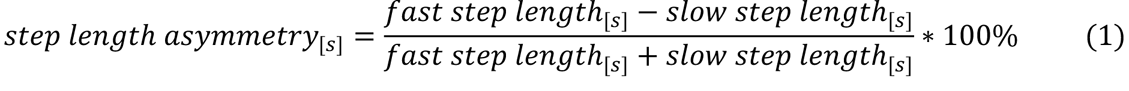

Thus, values of 0 indicate perfect symmetry between the fast and slow step lengths, while values further from 0 indicate greater asymmetry. We baseline corrected this measure by subtracting the mean step length asymmetry during the Baseline phase from each stride in the experiment for each individual. We removed outlier step length asymmetry strides, defined as any step length asymmetry exceeding 3x the interquartile range of that participant’s step length asymmetry (mean percent removed [min max] stroke = 0.14% [0.0 0.39]; control = 0.92% [0.0 2.32]). This prevented strides outside the normal range of variability from influencing the results. We also removed the first stride of each phase to account for treadmill acceleration.

We used step length asymmetry data to calculate an Adaptation Index for each individual for each stride (s; Charalambous et al. 2018; Mawase et al. 2014; Roemmich et al. 2016):

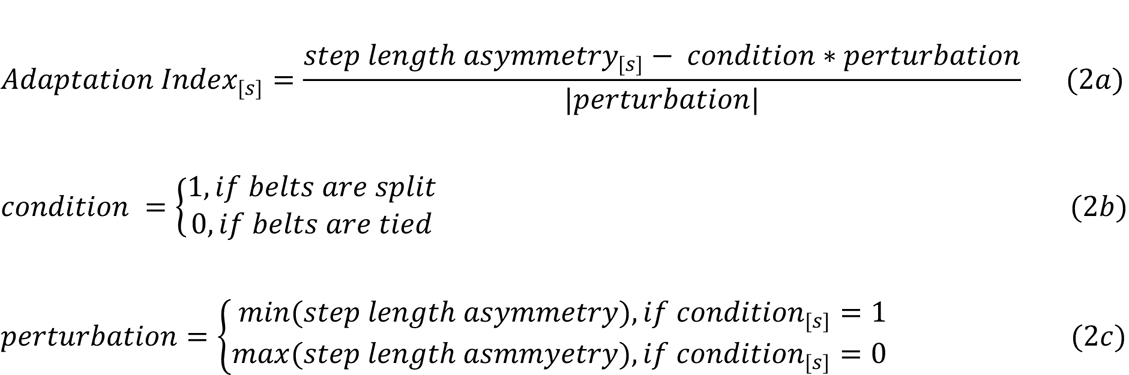

The min and max step length asymmetries used to determine the perturbation in equation 2c were calculated only within the first 10 strides of each respective phase (Charalambous et al. 2018). Thus, during the Adaptation phase, an Adaptation Index of 0 represents the minimum step length asymmetry (i.e., the max perturbation), and 1 indicates the perturbation has been fully corrected. The reverse is true during the De-adaptation phase.

To test our hypotheses, we averaged Adaptation Index during 4 key timepoints of interest: 1) Feedback On: the final five strides of the feedback being on during Adaptation, 2) Feedback Off: the first five strides immediately after the feedback was turned off during Adaptation, 3) End Adaptation: the last five strides of Adaptation, and 4) Implicit Aftereffect: the first five strides of De-adaptation. Our hypothesis for explicit learning was tested by comparing Feedback On (implicit adaptation plus explicit learning), to Feedback Off (implicit adaptation only). Larger differences between Feedback On and Feedback Off indicate greater explicit learning. We assessed the interaction between group (stroke vs control) and time (Feedback On vs Feedback Off) as our primary behavioral measure testing for impaired explicit learning in the stroke group. A secondary behavioral measure of explicit learning was between group differences during Feedback On, as this reflected the ability of individuals to use the visual feedback during implicit adaptation. Our primary behavioral measure to assess implicit adaptation, defined as the recalibration of step lengths caused by the split-belt treadmill, was comparing the Implicit Aftereffect between groups (Reisman et al. 2005; Shadmehr and Mussa-Ivaldi 1994). A secondary behavioral measure of implicit adaptation was a between groups comparison at End Adaptation.

### Computational modeling

Since the behavioral analysis only provides a brief (i.e., 5 strides) and somewhat arbitrary window into explicit learning and implicit adaptation, we used a computational model to characterize each learning process. This approach allowed us to map the underlying learning processes and their subcomponents onto each participant’s behavior. Specifically, we fit the model to individual data to obtain a unique set of parameter values for each participant. Since these parameters represent specific aspects of explicit and implicit learning, we can make inferences regarding the function of these underlying learning components (Charalambous et al. 2018; Mawase et al. 2014; Roemmich et al. 2016). Then, we compared the individual learning processes (i.e., model parameters) between the stroke and control groups.

This “voluntary correction” model was previously used to capture explicit and implicit learning in this paradigm (Roemmich et al. 2016), and the implicit adaptation component of the model can successfully capture split-belt adaptation behavior in individuals with stroke (Charalambous et al. 2018). The computational modeling used here followed that of Roemmich and colleagues which defines the Adaptation Index (x) on each stride (s) as the sum of both explicit learning (e_explicit_) and implicit adaptation (x_implict_):

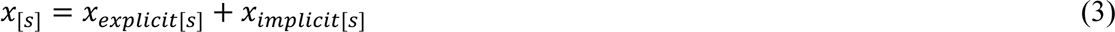

Both processes correct for the same error (error_[s]_ = perturbation_[s]_ − x_implict[s]_), where the perturbation = 1 during Adaptation and 0 during De-adaptation. Therefore, this model assumes that the nervous system is correcting for this error using the implicit processes, and it further reduces the error using the explicit process (Roemmich et al. 2016). Explicit learning is only active when the feedback is on:

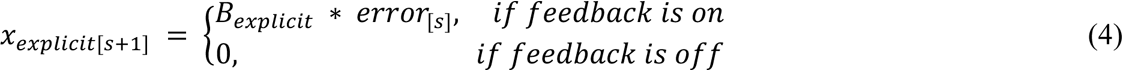

The free parameter, B_explicit_, represents the learning rate for explicit learning as it is the proportion of error that is explicitly corrected from one stride to the next (i.e., higher values indicate faster learning). The implicit adaptation process has dual components, fast and slow, and is active throughout the Adaptation and De-adaptation phases (Roemmich et al. 2016; Smith et al. 2006):

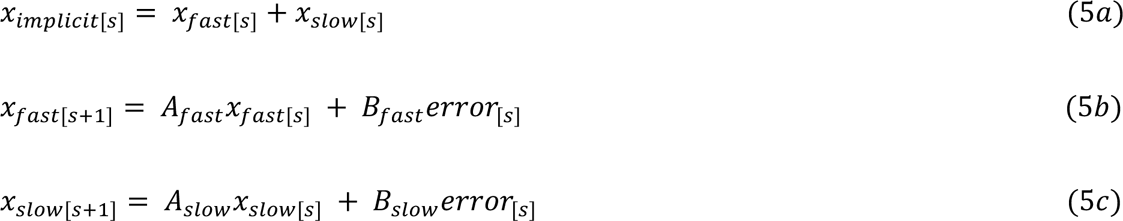

Implicit learning has four free parameters. The learning rates, B_fast_ and B_slow_, represent the proportion of the error that is implicitly corrected from one stride to the next, and the retention rates, A_fast_ and A_slow_, represent the proportion of the current adapted state that is retained. The fast process quickly learns from errors, but also quickly forgets, while the slow process takes longer to learn from errors but retains longer (Smith et al. 2006).

### Model fitting and model comparison

We fit the voluntary correction model to each participant’s Adaptation Index data during the Adaptation and De-adaptation phases using MATLAB’s fmincon function, setting the objective function as the sum of squared errors between the model output (x) and the data. All parameters were constrained between 0 and 1. Additionally, we constrained the fast- learning rate, B_fast_, to be at least 5 times higher than the slow learning rate, B_slow_; and the slow retention rate, A_slow_, was constrained to be greater than the fast retention rate, A_fast_ (Roemmich et al. 2016; Smith et al. 2006). To ensure stable fits, we initialized the implicit process parameters to the same values (Charalambous et al. 2018; Roemmich et al. 2016; Smith et al. 2006) based on a prior locomotor adaptation study in individuals with stroke (Charalambous et al. 2018): A_fast_ = 0.92, B_fast_ = 0.03, A_slow_ = 0.996, B_slow_ = 0.004. The explicit parameter was initialized at uniformly random values between 0 and 1. To improve the stability of the B_explicit_ parameter, we performed 10 initializations for each participant. We calculated individual fits (r^2^) to this model, but due to the greater stride-to-stride variability and idiosyncratic behavior of some individuals with stroke, the fits were worse for the stroke group compared to controls. As we were primarily interested in capturing group-level behaviors, we also performed a bootstrapping procedure to obtain a more consistent estimate of both group’s fits, while also quantifying our level of uncertainty around the sample means (Charalambous et al. 2018; Kim et al. 2019; Smith et al. 2006; Taylor and Ivry 2011; Tsay et al. 2021). We resampled with replacement the stride-by-stride data across participants 1000 times separately for each group, obtaining an average learning function for each bootstrap sample. We then fit the model and calculated an r^2^ value for each bootstrapped sample. We report the mean and 95% confidence intervals of these r^2^ values.

To ensure that we did not overfit the data with the five-parameter, voluntary correction model, we also fit two simpler models to the data. For the single-rate model, the motor output (x; i.e., Adaptation Index) is the result of a single process that has two parameters B, the learning rate, and A the retention rate (Smith et al. 2006; Thoroughman and Shadmehr 2000):

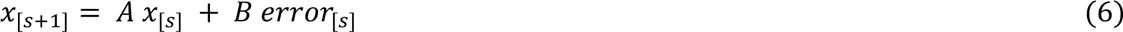

Additionally, we fit a four-parameter, dual-rate model to the data (Charalambous et al. 2018; Roemmich et al. 2016; Smith et al. 2006). This dual-rate model represents motor output (x; i.e., Adaptation Index) as the sum of a fast and slow process. This model is equivalent to the implicit process in the voluntary correction model (equations 5a-c). Thus, neither alternative model includes a voluntary correction, or explicit learning component. We compared fits of the three different models using Akaike Information Criterion (AIC). To determine the method of model comparison, we performed a model recovery analysis. We simulated the experiment 100 times with each model, the subsequently fitted each model to the three simulations. We calculated the number of times the model that simulated the data was best at fitting its own simulation (which should be expected if the models can be distinguished; Wilson and Collins 2019). We performed this procedure using both AIC and BIC and found that AIC was better at distinguishing between the three models for the current experiment (Wilson and Collins 2019).

### Statistical analysis

For our Bayesian statistical analysis, we assumed the Adaptation Index data at each timepoint of interest (Feedback On, Feedback Off, End Adaptation, Implicit Aftereffect) was sampled from a normal distribution, with a mean which depended on both within subject (i.e., time) and between subject (i.e., group) parameters as well as an interaction parameter. We confirmed a normal distribution was a reasonable assumption for our data by observing favorable model comparison results against a student’s t distribution. We estimated the posterior distribution for these means (and all the statistical model parameters) using Bayes rule, combining the evidence from our data with our prior assumptions about each parameter. The prior distributions for the between and within subject effects were set as a wide normal distribution centred on 0, and the prior for the standard deviation was set as a wide, positive-valued uniform distribution. Thus, our prior assumptions only served to make our inferences more conservative and did not bias the posterior. We estimated the joint posterior distribution in Python v4.3.0 using the PyMc 4 library (Salvatier et al. 2016) and the bayes-toolbox Python package (Kim 2023). We used Markov Chain Monte Carlo sampling to sample from joint posterior distribution 10,000 times with 2,000 tuning samples. We performed posterior predictive checks to ensure that the posterior samples accurately represented the data (Kruschke 2014; McElreath 2016).

This procedure allowed us to report the full range of credible differences between the groups along with the probability of a difference, given our data. To accomplish this, we compared the posterior distributions of between group differences which are presented as histograms representing the full distribution of possible differences based on the data we collected. For each posterior distribution, we report the mean and 95% high density interval (HDI), defined as the narrowest span of credible values that contain 95% of the distribution (Kruschke 2014). The HDI can be interpreted as the true value falling between this range with 95% certainty. Therefore, the 95% HDI provides an estimate of the size of the effect with 95% certainty. We also report the probability of a difference as a percentage of posterior distribution samples on one side of zero. Therefore, the p_difference_ value provides an estimate of the reliability of the effect. We do not provide a specific decision rule for the p_difference_ values as we would an alpha value. Rather, we made our inferences based on the actual degree of certainty of the difference, with larger p_difference_ values indicating greater certainty.

## Results

Of the 21 individuals recruited to participate in the stroke group, we removed 4 from the analysis because they either could not complete the task (1 due to double vision, and 2 due to fatigue that required stopping the treadmill) or they did not follow instructions (n=1). Average participant characteristics for each group are displayed in Table 1. In Figure 2, we display the mean, baseline-corrected step length asymmetry data during the Adaptation and De-adaptation phases for both groups. For ease of group comparisons, we present our primary analyses using the Adaptation Index. We note that similar results were obtained when using step length asymmetry index, with no impact on any of our inferences.

**Figure 2.**
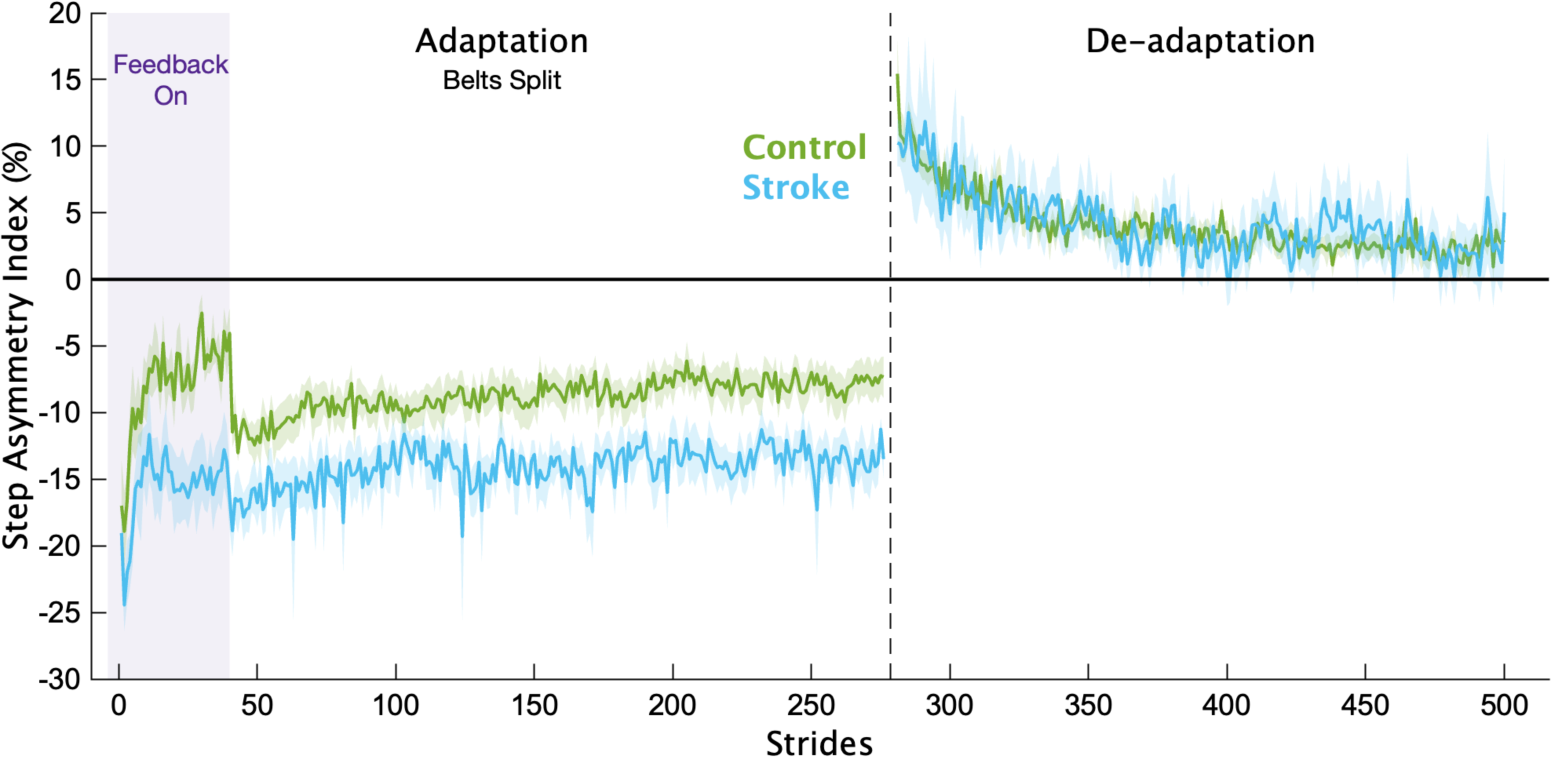
Step length asymmetry. Mean baseline-corrected step length asymmetry for each group for the Adaptation and De-adaptation phases. Purple shading is the time when the feedback was on. The vertical dashed line separates the Adaptation and De-adaptation phases. Each phase was truncated to the participant with the shortest phase for visualization purposes. Shading represents SEM.

**Table 1.** Group characteristics. Demographic and clinical characteristics of participants. All continuous variables are represented as mean ± 1 SD. (F = female, M = male, R = right, L = left)

|  | Stroke Group<br>(n=17) | Control Group<br>(n=18) |
| --- | --- | --- |
| Age (years) | 64.5 ± 10.2 | 64.8 ± 9.6 |
| Sex | 9M / 8F | 9M / 9F |
| Time since stroke (months) | 71.0 ± 49.1 |  |
| Side of brain lesion | 7R/10L |  |
| Self-selected (overground) walking speed (m/s) | 0.92 ± 0.27 | 1.33 ± 0.28 |
| Fastest (overground) walking speed | 1.29 ± 0.38 | 1.81 ± 0.22 |
| Fast treadmill belt speed | 0.94 ± 0.12 | 1.00 ± 0 |
| Lower Extremity Fugl Meyer | 25.41 ± 6.27 |  |

In Figure 3, we display the mean Adaptation Index data and key timepoints of interest for each group. First, we determined if individuals with stroke had impairments in explicit learning (Figure 3B). Based on our instructions and previous work (Roemmich et al. 2016), we assumed participants used explicit learning only while the feedback was on during the Adaptation phase. Therefore, explicit learning magnitude was characterized as the difference in Adaptation Index between Feedback On and Feedback Off. This difference was larger for the control group (mean interaction effect [95% HDI] = 0.09 [-0.05 0.25], p_difference_ = 88.2%), providing evidence that the individuals with stroke had diminished explicit learning compared to controls. Additionally, individuals in the stroke group were less able to use the visual feedback during Adaptation compared to controls (Figure 3C), with much lower Adaptation Index values during Feedback On (mean group difference = 0.23 [0.11 0.34], p_difference_ = 100.0%). Combined, these results point to impairments in explicit learning in individuals with stroke compared to controls.

**Figure 3.**
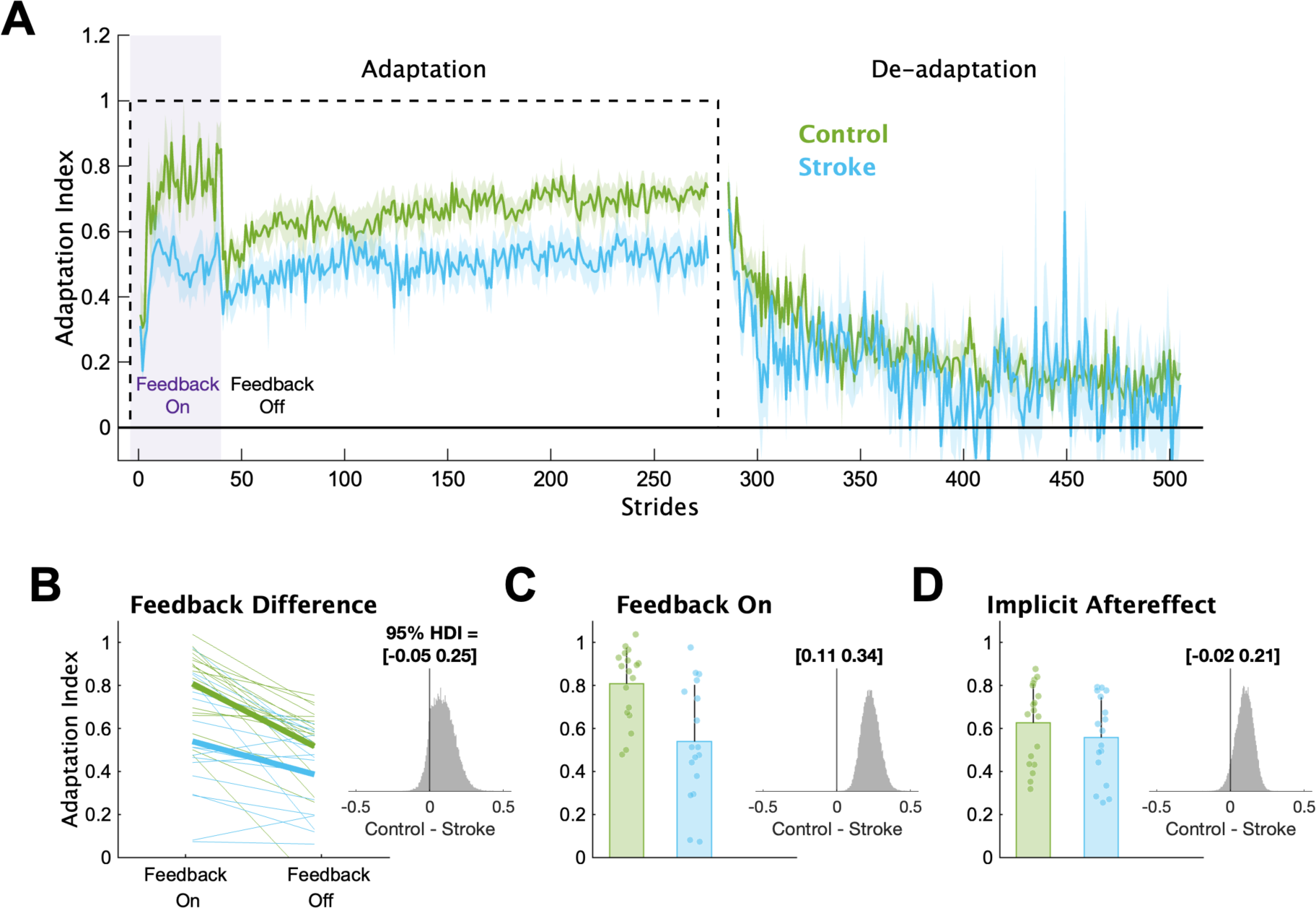
Adaptation Index. (**A**) Group averaged Adaptation Index data for the Adaptation and De-adaptation phases. The dashed line represents the walking period when the belts were split (i.e., the perturbation). Purple shading represents the time when the feedback was turned on. For visualization purposes, data for each phase were truncated to the individual with least number of strides. Solid lines represent group means, and shading represents SEM. (**B**) Group and individual data for the Feedback On and Feedback Off timepoints. Thick lines represent the group average slopes. **(C)** Group and individual data for the Feedback On timepoint. **(D)** Group and individual data for the Implicit Aftereffect timepoint. For panels C-D, bars represent group means, error bars represent 1 SD and smaller dots represent individuals. The insets display a histogram of the posterior distribution for the between group differences. The black vertical line in the histogram is there to aid visualization of the credibility of a between group difference (i.e., how much of the posterior probability distribution is on one side of zero). We report the 95% HDI regarding the range of credible effect sizes above the insets of the posterior distributions.

Next, we determined if individuals with stroke had impaired implicit adaptation by comparing the size of the implicit aftereffect (Figure 3D). The control group demonstrated larger implicit aftereffects compared to the stroke group (group difference = 0.10 [-0.02 0.21], p_difference_ = 94.3%), providing evidence that individuals with stroke have impaired implicit adaptation compared to controls. Additionally, we found large and reliable differences between the groups at End Adaptation (group difference = 0.17 [0.07 0.28], p_difference_ = 99.9%). Overall, the behavioral results indicate that impairments may exist in both explicit learning and implicit adaptation.

To provide a more complete picture of how each individual learning process contributed to overall adaptation, we applied a series of computational models to the data. The voluntary correction model, specifically, allowed us to map each individual’s behavior to explicit learning and implicit adaptation processes (Figure 4; see supplemental Figures S1-S2 for individual fits). This model fit the bootstrapped data well (stroke group mean r^2^ [bootstrapped 95% CI] = 0.70 [0.11 0.89]; control group = 0.90 [0.81 0.95]). The model fits to the individual stroke group’s data were lower on average as compared to the control group (mean [95% HDI] stroke = 0.31 [0.16 0.47]; control = 0.65 [0.53 0.77]), which is not surprising given that stroke participants tend to be more heterogeneous than age-matched controls. Nevertheless, based on the bootstrapping analysis (Fig. 4A and B), the voluntary correction model reasonably approximated the systematic group-level behavior caused by the experimental paradigm, as was our goal. Importantly, we confirmed that the voluntary correction model had better (lower) AICs then both the single rate and the dual rate model for both the stroke (single rate AIC difference mean [95% HDI] = 111 [54 167], p_difference_ = 100.0%; dual rate AIC difference = 36 [-20 89], p_difference_ = 89.9%) and control groups (single rate AIC difference = 281 [226 336], p_difference_ = 100.0%; dual rate AIC difference = 89 [35 142]; p_difference_ = 99.9%), indicating that the voluntary correction model accurately characterizes learning on this task without overfitting. As the single- and dual-rate state-space models do not include a voluntary correction process, these results also support our assumption that explicit learning contributed to behavioral change specifically when visual feedback was on. Overall, this analysis provides further support for quantifying the level of explicit learning impairment for the stroke group using the B_explicit_ parameter Comparing the individual parameters from the voluntary correction model allowed us to determine the specific components of learning that were impaired. The learning rate parameter for explicit learning, B_explicit_, served as a measure of each individual’s explicit learning ability, with higher values indicating faster explicit learning (Figure 4C). The stroke group had much smaller B_explicit_ values compared to the control group (group difference = 0.23 [0.06 0.40], p_difference_ = 99.5%), providing strong support for the hypothesis that explicit learning is impaired in individuals with stroke compared to controls. Next, we examined the four implicit adaptation process parameters (Figure 4F-G). While there was evidence of differences between groups for most parameters, the magnitude of differences for three of the four were near zero (group differences: A_slow_ = 0.00 [-0.00 0.01], p_difference_ = 87.7%; B_slow_ = 0.00 [-0.00 0.00], p_difference_ = 68.2%, B_fast_ = 0.01 [-0.01 0.03], p_difference_ = 84.7%). In contrast, there was a marked difference in the retention rate for the fast state (A_fast_ group difference = 0.09 [-0.04 0.24], p_difference_ = 92.6%). Thus, it appears that individuals with stroke, as a group, have a specific impairment in their ability to retain what was learned by the fast implicit adaptation process. In sum, the results of our computational modeling provided strong support for the hypothesis that explicit learning is impaired post-stroke and revealed that the retention rate for the fast state could underlie slower implicit adaptation in stroke.

**Figure 4.**
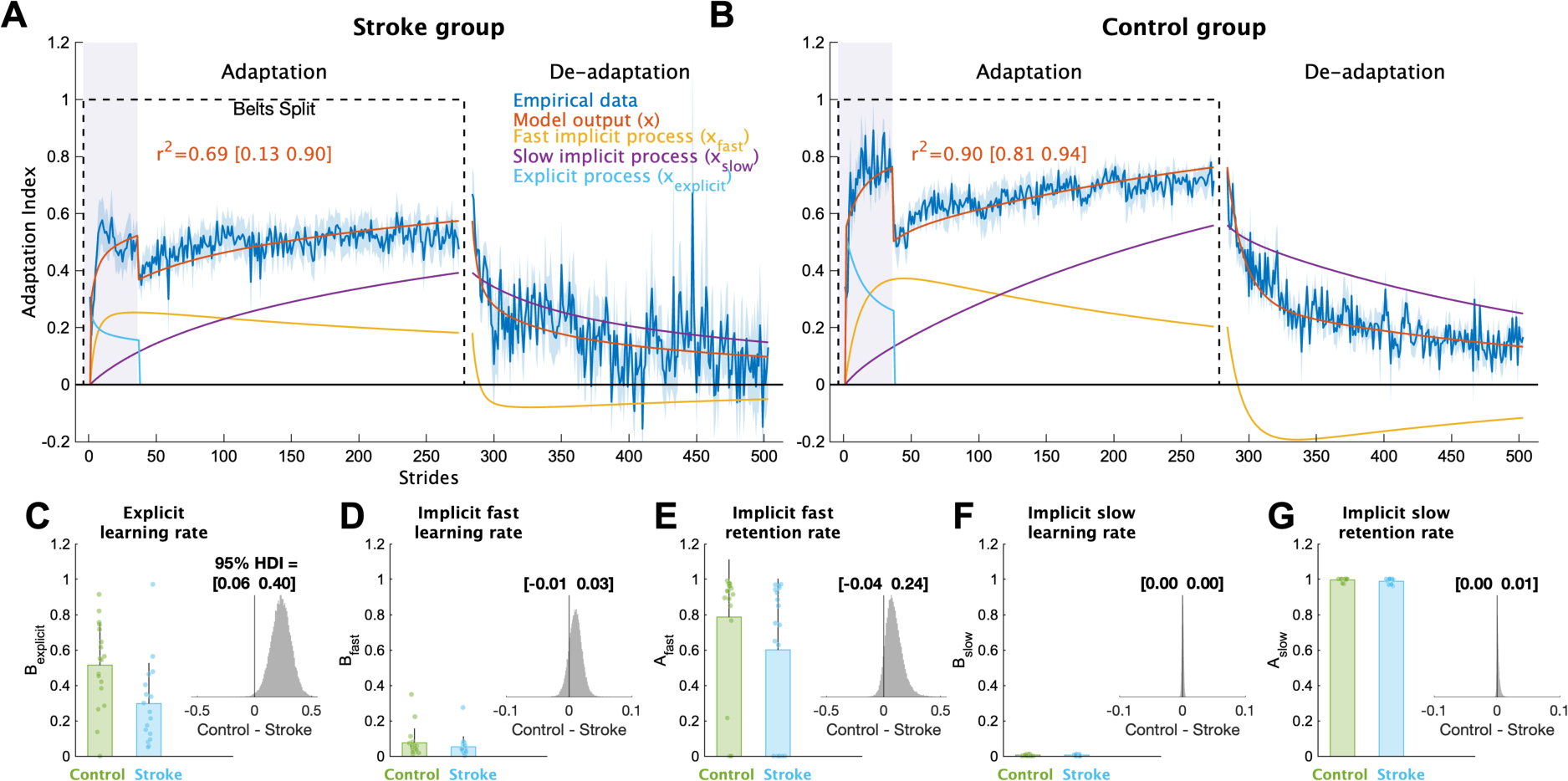
Computational model results. Mean model fits to bootstrapped samples plotted against the bootstrapped stride-by-stride data for the **(A)** stroke group and **(B)** control group. See supplemental Figures 1 and 2 for the model fits for each individual participant. Purple shading represents the time when the feedback was turned on. For visualization purposes, data for each phase were truncated to the individual with least number of strides. Shading represents 1 SD of the bootstrapped samples (i.e., the standard deviation of sample means or standard error of the mean) **(C-G)** Model parameter values for each group. Bars represent group means and error bars represent 1 SD and smaller dots represent individuals. The insets are histograms of the posterior of the between groups difference (contrast) in parameter values. We report the 95% HDI regarding the range of credible effect sizes above the insets of the posterior distributions. Note the scale of the x-axis varies for these inset plots.

## Discussion

In the current study, we examined explicit learning and implicit adaptation within the same locomotor learning task in individuals with chronic, hemiparetic stroke. We combined a behavioral manipulation and computational modeling to determine the presence, and potential degree, of impairment in both learning processes. The majority of work in locomotor learning and stroke has primarily studied implicit adaptation (Betschart et al. 2017; Charalambous et al. 2018; Reisman et al. 2007; Savin et al. 2013; Tyrell et al. 2014), with less attention paid to explicit learning (French et al. 2021b, 2021a). While some gait studies have examined both processes in the same task (Cherry-Allen et al. 2018; Day et al. 2019; Moore et al. 2022; Mutha et al. 2011; Schaefer et al. 2009), only two studies in reaching have attempted to discern their individual contributions to overall motor learning (Binyamin-Netser et al. 2023; Taylor and Ivry 2014). To our knowledge, the current study is the first to assess explicit and implicit motor learning within the same task in individuals with chronic stroke using both behavioral manipulations and computational modeling. While our sample size is relatively small, our results provide strong evidence that stroke impairs explicit learning and the rate of implicit adaptation during a locomotor task that elicits dissociable contributions from both processes. This has important implications for the design of locomotor learning tasks in post-stroke rehabilitation because many interventions involve both explicit and implicit components.

### Explicit learning is impaired in chronic stroke

We found that individuals with chronic, hemiparetic stroke have impairments in explicit learning in a locomotor task involving both explicit learning and implicit adaptation. Individuals with stroke had a smaller change in behavior compared to controls after the visual feedback, intended to drive and support explicit learning, was removed. Additionally, the computational modeling revealed significantly slower explicit learning in individuals with stroke. To ensure these results were not due to motor control deficits that prevented participants with stroke from effectively implementing explicit strategies, we performed a series of Bayesian regressions to test the effect of lower extremity function (quantified by Lower Extremity Fugl-Meyer [LEFM] scores) on our learning measures. Our analyses showed that there was essentially no effect of LEFM on the B_explicit_ parameter (beta coefficient = 0.005 [-0.01 0.02]), or on our behavioral measure of explicit learning (beta coefficient = 0.01 [-0.004 0.02]), providing further evidence that the behavioral differences observed were due to explicit learning deficits in participants with stroke.

While there are few prior studies assessing both explicit learning and implicit adaptation in the same task after stroke, one experiment in reaching showed that individuals with lateral prefrontal cortex (LPFC) lesions demonstrate impairments in explicit learning (Taylor and Ivry 2014). This and other work raises the possibility that cognitive processes such as working memory or general cognition in reaching studies (Anguera et al. 2011; Binyamin-Netser et al. 2023; Seidler et al. 2012; Vandevoorde and Orban de Xivry 2020), and fluid cognition in gait (French et al. 2021a) contribute to explicit motor learning, but more work is required to determine the specific contribution of cognition to explicit learning in stroke. Future work could use techniques such as lesion mapping along with computational modeling to provide a mechanistic explanation for the explicit learning impairments we observed.

Contrary to the current findings, prior work in reaching (Binyamin-Netser et al. 2023) and gait (French et al. 2021b, 2021a) observed no differences in explicit learning in individuals with stroke compared to controls. However, the studies in gait used a primarily explicit task without a split-belt perturbation, likely making the task easier, which could reduce the ability to detect explicit learning deficits in stroke. The reaching study dissociated explicit learning and implicit adaptation using visual cues (the color and shape of a cursor). It is possible that either this manner of distinguishing between explicit and implicit processes or the broader inclusion criteria for their stroke group can account for the differences between their findings and those of the current study. Similar to Taylor and Ivry (Taylor and Ivry 2014), we provided clear instructions and removed all visual feedback to ensure explicit learning was minimized, and provided a narrower range of inclusion criteria, potentially explaining why our results were more consistent with theirs. Still, it is critical to determine if the manner of eliciting explicit learning (a specific type of cue or instruction) impacts the ability to use this process in stroke given its ubiquity in rehabilitation settings.

### Slower implicit adaptation in stroke is due to worse retention of the fast process

Contrary to our hypothesis, we found evidence that implicit adaptation is impaired after stroke. The stroke group demonstrated smaller implicit aftereffects and a lower plateau at the end of the Adaptation phase indicating a smaller overall magnitude of implicit adaptation. Prior studies in locomotor adaptation after stroke indicate that the overall magnitude of implicit adaptation is similar to controls, but the rate is slower (Malone and Bastian 2014; Reisman et al. 2007; Savin et al. 2013; Tyrell et al. 2014). Therefore, the slower rate of adaptation in stroke, combined with the relatively short Adaptation phase in the current study (8 minutes compared to 10-15 minutes in the prior studies), could have prevented us from observing asymptotic adaptation. While we assume that these impairments in implicit adaptation during split-belt walking are related to learning from sensory prediction error, it is also possible that split-belt adaptation is driven by energy optimization (Finley et al. 2013; Sánchez et al. 2019, 2020; Sánchez and Finley 2018), a theory that could be tested in individuals with stroke.

While it may seem that the visual feedback interfered with implicit adaptation for the stroke group, prior work in young individuals with intact neurologic systems demonstrate that visual feedback used to either help or hinder performance during split-belt walking does not change the total magnitude of implicit adaptation (Long et al. 2016; Malone and Bastian 2010; Roemmich et al. 2016). Additionally, individuals with stroke can successfully adapt to the split-belt treadmill while also explicitly learning to change a separate gait parameter (knee flexion angle) using visual feedback (Cherry-Allen et al. 2018). Therefore, it is unlikely that explicit learning itself hindered implicit adaptation in the current study since implicit adaptation proceeds in spite of explicit learning, and even worsens performance in some cases (Mazzoni and Krakauer 2006; Taylor and Ivry 2011), across reaching and walking paradigms (Mazzoni and Krakauer 2006; Roemmich et al. 2016), including in stroke (Binyamin-Netser et al. 2023; Cherry-Allen et al. 2018; Taylor and Ivry 2014).

The computational modeling utilized in this study provides insight into why the learning rate of implicit adaptation was impaired in stroke in this task. The voluntary correction model incorporates a dual-rate model of adaptation which frames implicit adaptation as the combination of a fast state and a slow state (Smith et al. 2006). These states represent updates to an internal model, a prediction of the sensory consequences of movement, that could occur either in the cerebellum or motor cortex (Smith et al. 2006). One theory suggests that the motor cortex is responsible for retention of the adapted state while the cerebellum is responsible for learning (Galea et al. 2011; Li et al. 2001). Thus, damage to motor cortices or possibly its outputs could explain poor retention of the fast process in individuals with stroke. Alternatively, the fast process has been closely linked to explicit learning during visuomotor rotation tasks (Krakauer et al. 2019; Taylor and Ivry 2014). However, to date there is no evidence of contributions from explicit learning to standard split-belt adaptation (i.e., without additional visual feedback; Charalambous et al. 2018; Mawase et al. 2014; Roemmich et al. 2016). Another possibility is that the fast state represents a reactive balance element that is sensitive to environmental changes (Mariscal et al. 2023). Future studies are required to dissociate between these potential explanations.

## Conclusion

Motor learning involves multiple processes, both explicit and implicit, that work together to improve overall task performance. We found that individuals with chronic stroke have impairments in explicit learning and implicit adaptation during a locomotor task that elicits dissociable contributions from both. These findings are important because of the potential application to post-stroke rehabilitation, which often combines different forms of learning in a single task. To improve outcomes, future work should determine how locomotor rehabilitation interventions can be structured to target these deficits and optimize overall motor learning.

## Supplemental Material

Supplemental Figures S1–S2: https://osf.io/pws2k/.

## Data Availability

Source data for this study are openly available at https://osf.io/pws2k/

## Acknowledgements

We would like to thank Saunders Penn for helping to create Figure 1A.

A preprint of this manuscript is available at https://doi.org/10.1101/2024.02.04.578807.

## Author Contributions

JMW, DSR, HEK Conceived and designed research; HW, LF, JMW performed experiments; JMW analyzed data; JMW, ET, SMM, DSR, HEK interpreted results of experiments; JMW prepared figures; JMW drafted manuscript; All authors edited and revised manuscript; All authors approved final version of manuscript.

## Supplemental material

**Supplemental Figure S1.**
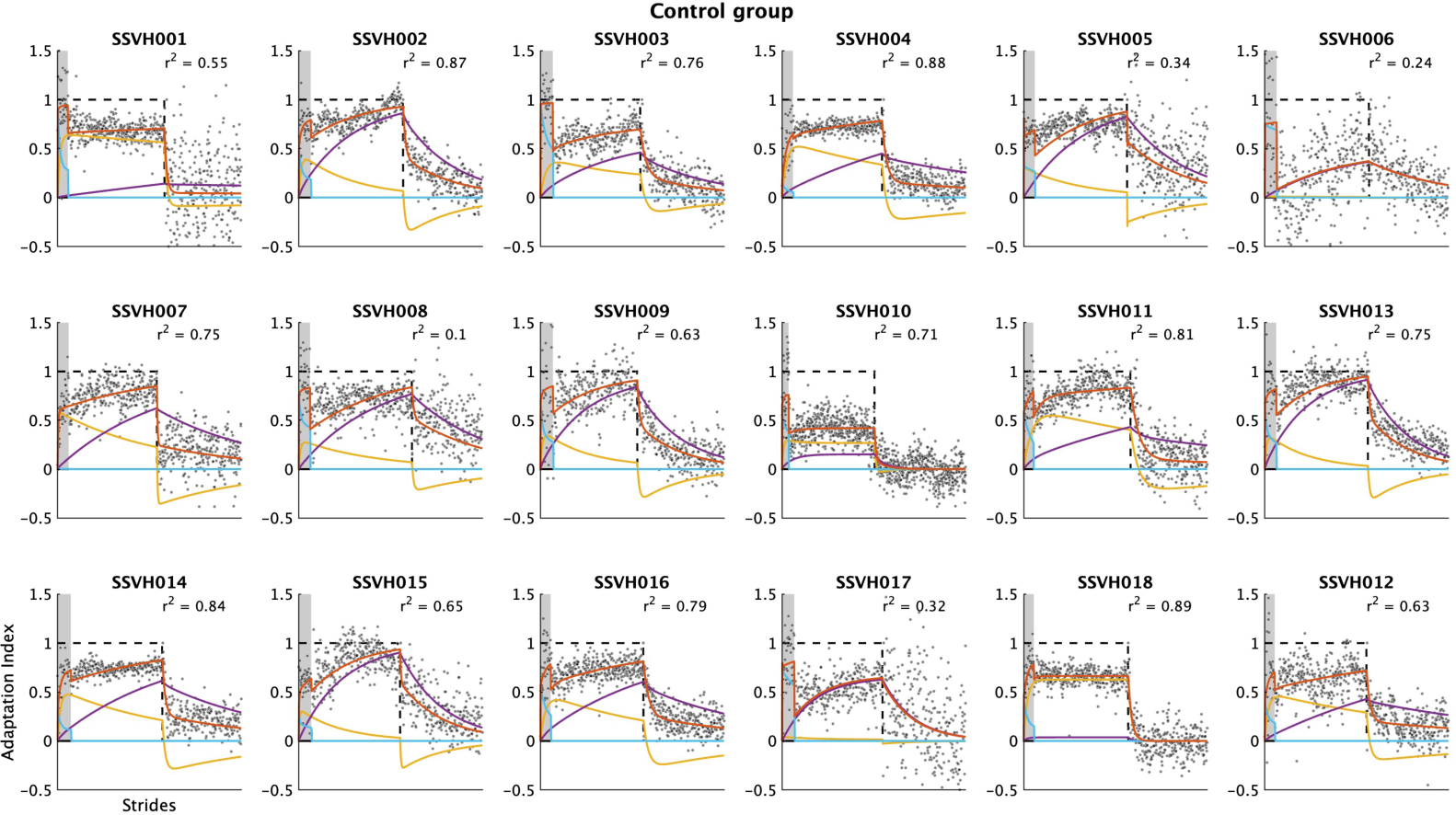
Individual model fits for the control group. The voluntary correction model was fit to each participant’s Adaptation Index data in the control group (n=18). Black dots represent the participant’s Adaptation Index on that stride and the black dashed line represents the perturbation (which equals one during the Adaptation phase and 0 during De-adaptation phase). The red function represents the model’s motor output (x), the light blue function (active only when the feedback is turned on – denoted by the gray shading) represents the explicit component (x_explicit_), and the yellow and purple functions represent the fast (x_fast_) and slow (x_slow_) components of the implicit process, respectively.

**Supplemental Figure S2.**
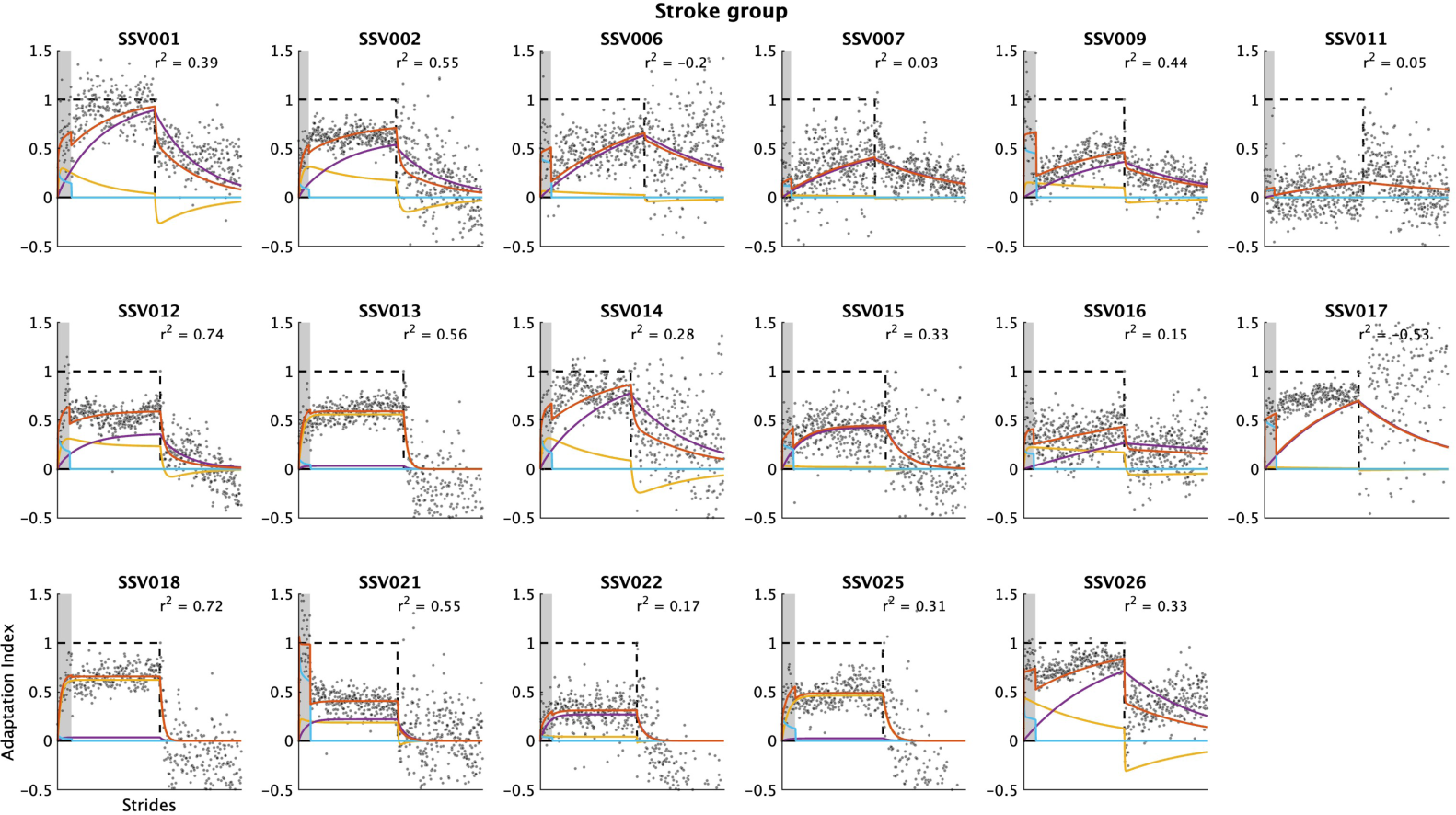
Individual model fits for the stroke group. The voluntary correction model was fit to each participant’s Adaptation Index data in the stroke group (n=17). Black dots represent the participant’s Adaptation Index on that stride and the black dashed line represents the perturbation (which equals one during the Adaptation phase and 0 during De-adaptation phase). The red function represents the model’s motor output (x), the light blue function (active only when the feedback is turned on – denoted by the gray shading) represents the explicit component (x_explicit_), and the yellow and purple functions represent the fast (x_fast_) and slow (x_slow_) components of the implicit process, respectively.

